# Instantaneous Beta Frequency Regulates Self-Generated Timing in Humans

**DOI:** 10.64898/2026.03.10.710730

**Authors:** Yitong Zeng, Xiangshu Hu, Zhenyi Hu, Jia Li, Qianyi Wu, Lu Shen, Biao Han

## Abstract

Self-generated timing requires the brain to determine when to terminate an ongoing action without reference to external cues. Although beta-band oscillations have been implicated in temporal control, how their intrinsic dynamics shape moment-to-moment variability in internally generated time remains unclear. Here, combining scalp EEG with intracranial recordings in humans, we show that the instantaneous frequency of beta oscillations (13-30 Hz) provides a dynamic control signal for self-generated timing. During a self-paced time production task, higher instantaneous beta frequency reliably predicted longer produced durations, both within individuals on a trial-by-trial basis and across individuals. This relationship was beta-specific, frontoparietally distributed, robust to motor and spectral confounds, and validated by convergent intracranial recordings. These findings run counter to classical pacemaker-accumulator models and instead establish beta-frequency dynamics as a mechanism that delays action termination by increasing the density of intrinsic stabilizing control updates during self-generated timing.

## Introduction

Accurate timing is a prerequisite for interacting with a dynamic world. While external sensory cues can entrain behavioral rhythms, humans possess a remarkable capacity to generate precise timing internally—a faculty essential for speech, music, and voluntary action(Merchant et al., 2013; Paton & Buonomano, 2018). In the absence of external references, the brain must construct and sustain an internal representation of elapsed time. Despite decades of research emphasizing ramping activity(Simen et al., 2011) or population-level dynamics(Paton & Buonomano, 2018), the neural mechanisms that account for the moment-to-moment variability of self-generated timing remain poorly understood.

Converging work suggests that neural oscillatory activity contributes to internally guided temporal control, with different frequency bands implicated across distinct neural systems(Frisoni et al., 2025; Milton & Pleydell-Pearce, 2016; Mioni et al., 2020; Morrow et al., 2024; Tsao et al., 2022; Wutz, 2024), including beta-band activity (13-30 Hz) within sensorimotor networks(Feurra et al., 2011; Kilavik et al., 2013; Kononowicz et al., 2018; Kononowicz & van Rijn, 2015; Kulashekhar et al., 2016; Wiener et al., 2018). In these networks, beta activity covaries with multiple aspects of temporal behavior: beta power is modulated in anticipation of predictable events(Fujioka et al., 2012; Kulashekhar et al., 2016) and scales with the duration of self-initiated movements(Bartolo et al., 2014). Beyond these correlational findings, causal interventions demonstrate that experimentally manipulating beta oscillations can bias motor timing(Schilberg et al., 2018; Wiener et al., 2018), motivating the idea that beta rhythms may provide a control signal for internally generated temporal goals. What remains unresolved, however, is how beta oscillations implement such temporal control on a moment-to-moment basis to bias the duration of ongoing actions.

Addressing this question requires moving beyond traditional descriptions of beta activity in terms of oscillatory power or phase. Such approaches implicitly treat beta rhythms as temporally stationary signals, an assumption that overlooks a fundamental property of neural oscillations: their intrinsic non-stationarity and transient fluctuations in instantaneous frequency(Cohen, 2014a). Consistent with this view, beta activity is increasingly recognized as time-varying, with spectral characteristics that evolve over behaviorally relevant timescales(Little et al., 2019; Sherman et al., 2016). Moreover, shifts in beta peak frequency are structured and behaviorally meaningful, tracking motor preparation and somatosensory processing independently of oscillatory power(Kilavik et al., 2013; Spitzer & Haegens, 2017). These observations motivate examining beta frequency at the single-trial, time-resolved level and raise the possibility that moment-to-moment beta-frequency dynamics may constitute a latent control parameter underlying internally generated timing.

Here, we directly test this possibility by asking whether the speed of the oscillation itself—the instantaneous beta frequency (IBF)—governs self-generated timing. Classical pacemaker-accumulator models predict that faster neural rhythms should function as faster clocks, leading to a more rapid accumulation of temporal “ticks” and consequently shorter produced durations(Gibbon, 1977; Treisman, 1963). In contrast, if beta oscillations serve to maintain the current sensorimotor state, as proposed by the “status quo” hypothesis(Engel & Fries, 2010), faster beta-frequency dynamics may increase the frequency of stabilizing control updates within a given time interval, thereby delaying motor state transitions and prolonging produced durations (Figure 1A). To adjudicate between these opposing accounts, we combined scalp electroencephalography (EEG) and intracranial stereo-EEG (sEEG) recordings while participants performed a self-paced time production task, estimating and producing a fixed time duration solely based on internal cues. By tracking instantaneous beta frequency across trials and individuals, this multimodal approach allowed us to determine whether beta-frequency dynamics provide a mechanistic link between intrinsic neural oscillations and self-generated temporal control.

**Figure 1.**
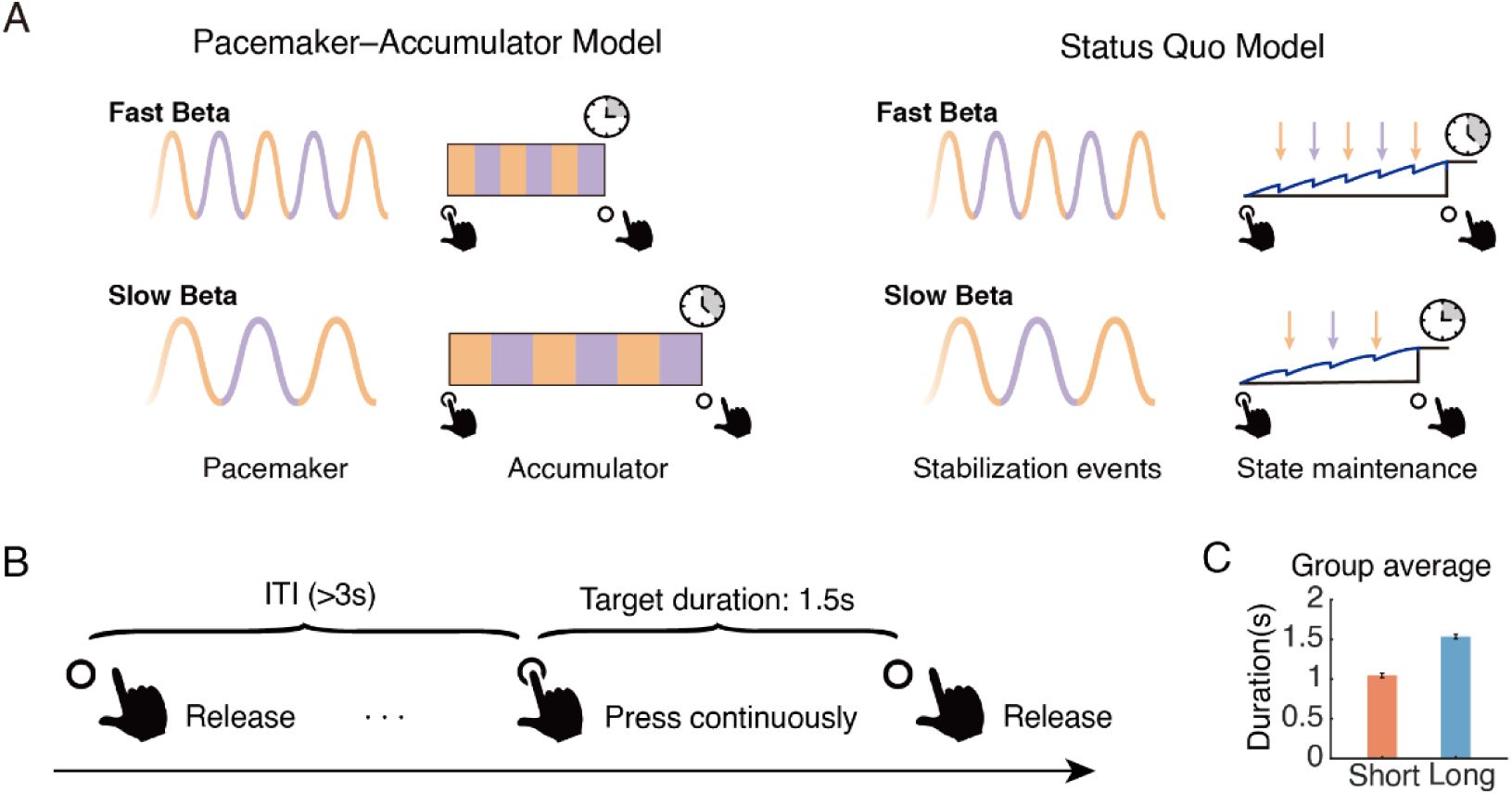
Conceptual frameworks, experimental paradigm, and behavioral results in the self-paced time production task. **A,** Left: Classical pacemaker–accumulator models assume that beta frequency determines the rate of temporal accumulation. Right: The status quo account proposes that beta frequency regulates the density of stabilizing control updates, thereby influencing motor state transitions and the timing of action termination. **B.** Schematic of a single trial in the self-paced time production task. Participants were instructed to press and hold a pressure sensor to internally produce a 1.5-s duration and release it when they judged the target duration had elapsed. A minimum gap of >3 s was required between consecutive keypresses. **C**, Mean self-timed durations for short and long trial tertiles averaged across participants. Error bars represent ±1 within-subject SEM.

## Results

We investigated how intrinsic neural dynamics support self-generated timing in the absence of external cues. Participants performed a self-paced time production task, in which they were instructed to press and hold a key to internally estimate a 1.5-s duration and then release it when they believed the target duration had elapsed (Figure 1B). Across participants, the overall mean produced duration was 1.28 ± 0.42 s, indicating a systematic underestimation of the objective 1.5-s target. A one-sample *t*-test confirmed that the produced durations were significantly shorter than the target duration (*t*_26_ = −2.787, *p* = 0.010). To examine within-subject variability, trials were sorted into tertiles based on each participant’s self-timed durations. The short-duration condition (lower tertile) had a mean of 1.04 ± 0.33 s, whereas the long-duration condition (upper tertile) averaged 1.53 ± 0.53 s (Figure 1C). As expected from the tertile-based grouping, durations in the long condition were longer than those in the short condition (*t*_26_ = 8.646, *p* < 0.001), confirming that the binning procedure effectively captured within-subject variability in produced time. Across participants, the mean short-duration values ranged from 0.77 to 1.97 s, whereas the mean long-duration values ranged from 0.91 to 2.62 s (Figure S1). These results demonstrate that subjective time production systematically diverged from the objective 1.5-s target and varied widely across individuals. Together, these behavioral results confirm that the task successfully elicited endogenous fluctuations in internal motor timing, providing a solid behavioral foundation for the subsequent neural analyses.

### Instantaneous beta frequency predicted self-timed duration

We first examined the overall relationship between oscillatory frequency dynamics and variability in self-generated timing across all scalp electrodes. Instantaneous frequency was computed at each electrode for the theta (4-8 Hz), alpha (8-13 Hz), beta (13-30 Hz), and gamma (30-120 Hz) bands, time-locked to key release. Among these frequency ranges, only the beta band exhibited a significant modulation between the long- and short-duration conditions (*cluster-based correction*, *p* = 0.012, significant cluster spanning −1.95 to −0.94 s relative to key release; Figure 2A). No reliable modulations were observed in the theta, alpha, or gamma bands, including when the gamma range was restricted to 30-50 Hz (*cluster-based correction*, all *p*s > 0.05; Figure S2A). To quantify this beta-band modulation, beta-frequency values were averaged across the pre-release period (−2 to 0 s) and compared between conditions. The instantaneous beta frequency (IBF), averaged across all electrodes, was significantly higher during long-duration than short-duration trials (*t_26_* = 3.16, *p* = 0.004; Figure 2B). Together, these results indicate that, within individuals, higher beta-band frequency during the pre-release period accompanies the production of longer self-timed durations.

**Figure 2.**
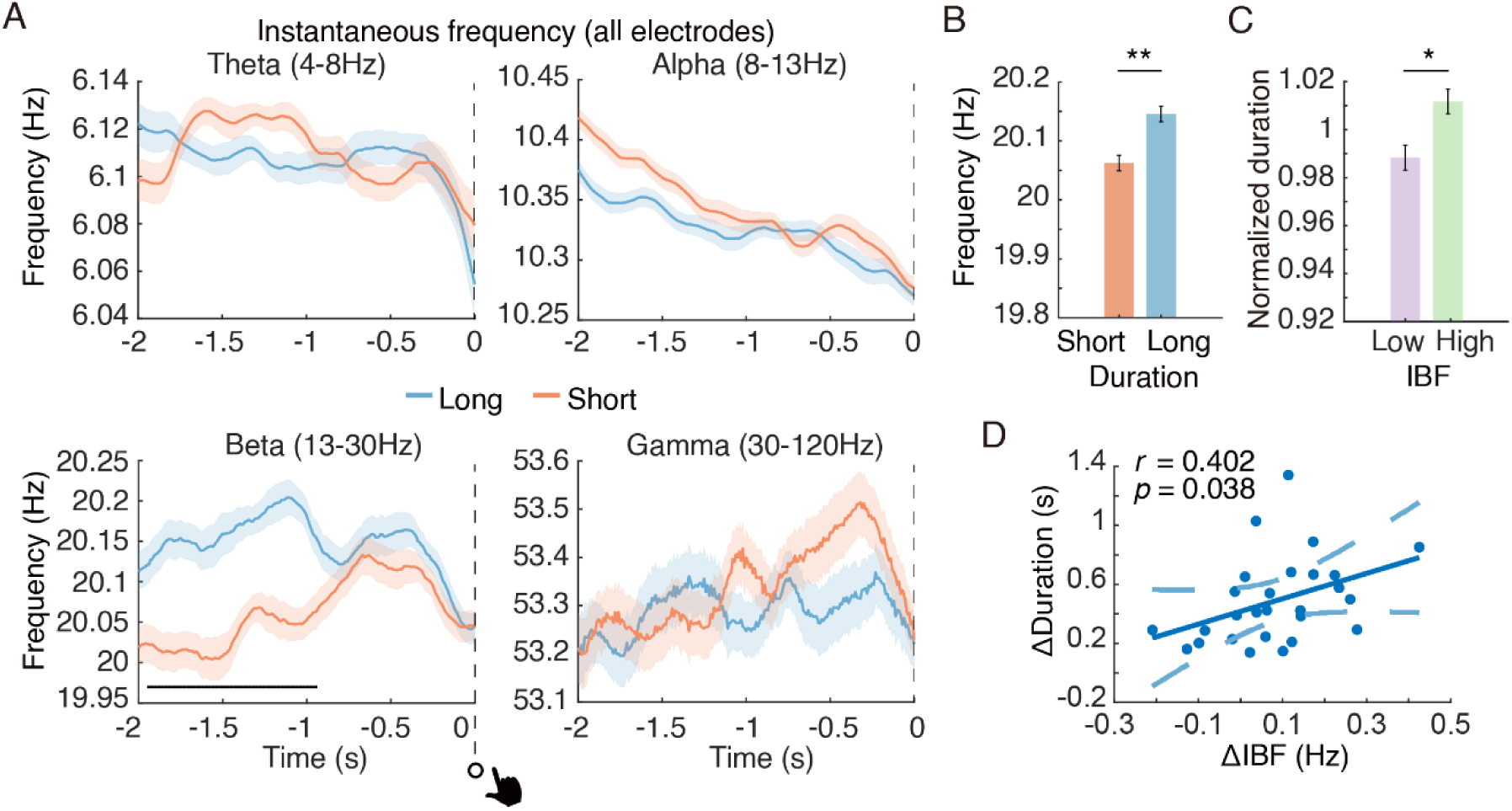
Relationship between the instantaneous frequency and self-timed duration in key-release-locked analysis. **A**, Within-subject analysis of the instantaneous frequency in the theta (4-8Hz), alpha (8-13Hz), beta (13-30Hz), and gamma (30-120Hz) bands. Significant time points are indicated by the horizontal black bar (*cluster-based correction*, *p* < 0.05). Shaded regions denote ±1 within-subject SEM. **B**, IBF values averaged across all electrodes during the pre-release period (−2 to 0 s) for the long- and short-duration conditions. **C**, Normalized self-timed durations for high- and low-IBF trials, with IBF computed across all electrodes during the pre-release period. Asterisks denote significant differences between conditions (\**p* < 0.05, ** *p* < 0.01). **D**, Correlation between the beta-frequency difference (long-short) averaged across all electrodes and the corresponding difference in produced durations across participants.

To determine whether this effect depended on the use of instantaneous-frequency estimates, we performed a complementary analysis using a conventional frequency-domain measure of beta oscillations. Peak beta frequency (PBF), estimated from FFT-based power spectra computed across all electrodes over the same pre-release window (−2 to 0 s), also differed significantly between long- and short-duration trials, with higher beta peak frequencies observed during long-duration trials (*t_26_*= 2.25, *p* = 0.033; Figure S3A). This convergence across analytic approaches indicates that the observed beta-frequency modulation is not specific to the instantaneous-frequency method.

We next assessed whether this relationship extended to trial-by-trial variability within individuals. Trials were divided into tertiles based on their IBF magnitude computed across all electrodes during the pre-release period (−2 to 0 s). Self-timed durations in the high-IBF trials were significantly longer than those in the low-IBF trials (*t_26_*= 2.26, *p* = 0.033; Figure 2C). This result indicates that moment-to-moment fluctuations in beta frequency within an individual predict trial-by-trial variability in the duration they produce, linking transient beta dynamics to fine-grained adjustments of internal timing.

At the between-subject level, this relationship generalized across participants: individuals who exhibited larger IBF differences (long - short) between the long and short conditions also produced larger differences in duration (long - short) (*r* = 0.402, *p* = 0.038; Figure 2D). This cross-participant correlation demonstrates that the strength of beta-frequency modulation scales with subjective time expansion across individuals, identifying a consistent neural signature of internal timing at the population level.

Finally, we tested whether the observed beta-duration relationship could be explained by differences in temporal precision rather than systematic shifts in produced durations. Because longer produced durations were, on average, closer to the target duration (1.5 s), an alternative interpretation is that higher frequency reflects more precise temporal estimation(Frisoni et al., 2025). To directly address this possibility, trials were ranked according to their absolute deviation from the target duration and divided into tertiles. Instantaneous beta frequency did not differ between high-precision and low-precision trials (*cluster-based correction*, all *p*s > 0.1; Figure S4A). The same null result was observed at the between-subject level: participants’ mean IBF was not correlated with their absolute deviation from the target duration (*r* = 0.020, *p* = 0.923; Figure S4B). Together, these control analyses demonstrate that beta-frequency dynamics are dissociable from temporal precision and instead systematically bias the duration of self-generated actions.

### Frontoparietal regions exhibited the strongest beta-frequency modulation

Building on the whole-scalp findings, we next mapped the spatial distribution of the beta-frequency modulation associated with self-timed duration. Beta-frequency values were averaged for each electrode within the pre-release period (−2 to 0 s relative to key release) for both the long- and short-duration conditions, and paired-samples t-tests were performed at the participant level. Electrodes exhibiting significant differences were marked as bold circular symbols on the scalp map. This analysis revealed a significant cluster over the frontal and parietal regions (*cluster-based correction*, *p* = 0.004; Figure 3A), indicating that the previously observed global modulation was driven predominantly by frontoparietal electrodes.

**Figure 3.**
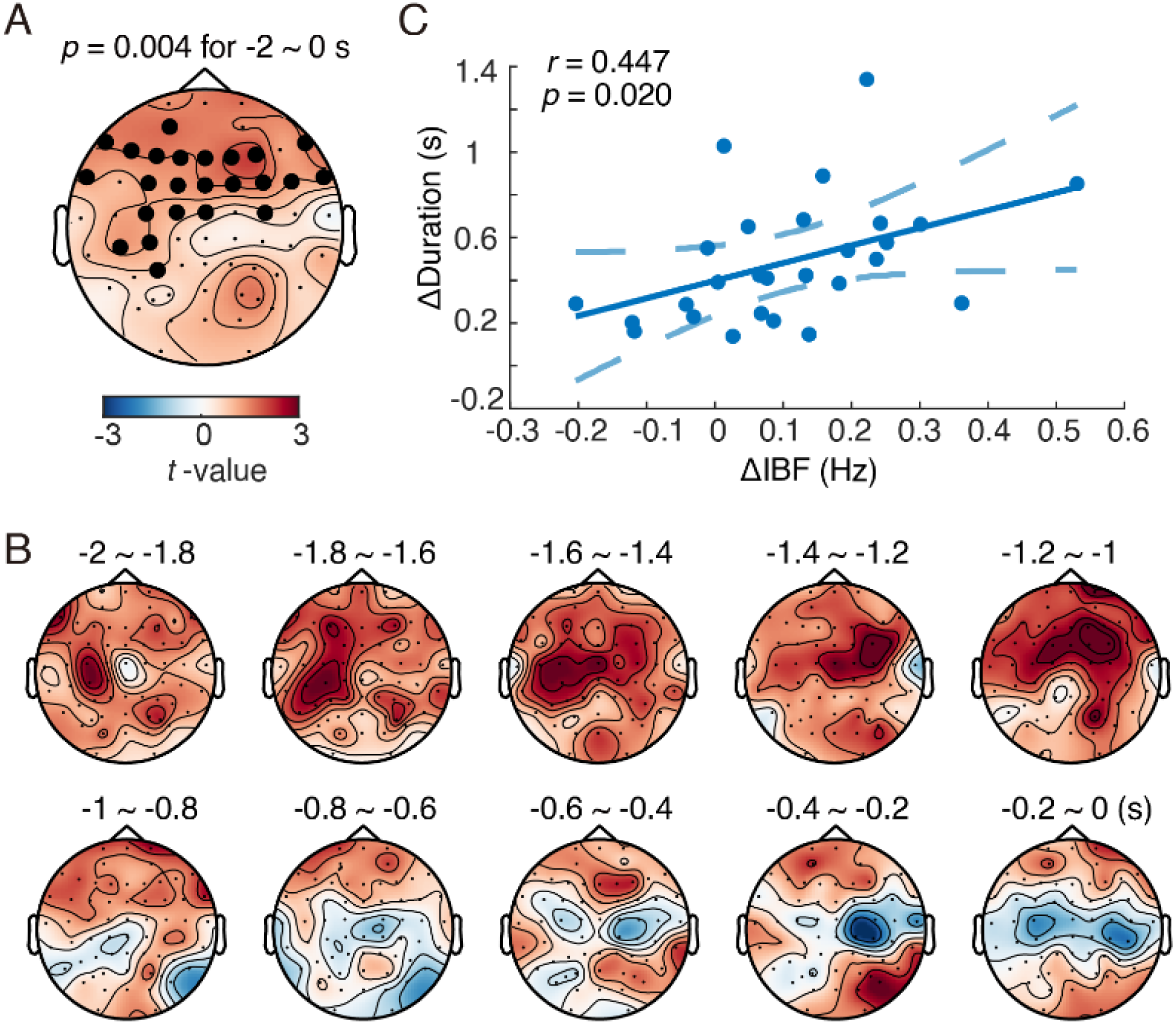
Spatial characterization of beta-frequency modulation. **A,** Scalp topography of beta-frequency modulation associated with self-timed duration. Beta-frequency values were averaged for each electrode within the pre-release period (−2 to 0 s relative to key release) for the long- and short-duration conditions. Electrodes showing significant differences (long > short, cluster-based corrected) are indicated by bold circular symbols. **B,** Dynamic scalp maps showing beta-frequency differences (long-short) across consecutive 0.2-s durations within the pre-release period, illustrating how the modulation evolved over time. **C,** Correlation between the beta-frequency difference (long-short) averaged across the significant frontoparietal electrodes identified in (A) and the corresponding difference in produced durations across participants.

To examine how this effect evolved over time, we computed dynamic topographies of beta-frequency differences between the long and short conditions across consecutive 0.2-s durations within the pre-release period. The frontoparietal enhancement emerged early and persisted throughout this duration (Figure 3B), reflecting sustained beta-frequency modulation as participants internally timed their key release.

We then assessed the behavioral relevance of this spatially localized modulation. The beta-frequency difference (long-short), averaged across the significant frontoparietal electrodes, correlated positively with the corresponding difference in produced durations (*r* = 0.447, *p* = 0.020; Figure 3C). Participants who exhibited stronger frontoparietal beta modulation produced greater self-timed durations difference, underscoring the key role of frontoparietal beta dynamics in supporting internally generated timing.

### Beta-frequency modulation was independent of motor processes

Because the beta-frequency effect identified in the key-release-locked analysis occurred close to the time of key pressing, it might, in principle, reflect motor-related activity associated with key pressing rather than internal timing mechanisms. To evaluate this possibility, we first examined whether the effect was driven by movement-evoked activity. We therefore re-aligned the EEG data to keypress onset and recomputed the difference in instantaneous frequency between the long- and short-duration conditions. A significant beta-frequency difference was observed from −0.45 to 0.05 s relative to keypress onset (*cluster-based correction, p* = 0.028; Figure 4A), whereas no reliable effects were found in the theta, alpha, or broad gamma (30-120 Hz) bands. To verify that the null gamma result was not due to the wide bandwidth, we also examined the low-gamma range (30-50 Hz), which likewise showed no significant modulation (Figure S2B). The beta-frequency modulation persisted under this motor-aligned analysis and, critically, the modulation began several hundred milliseconds before keypress onset and extended briefly after the press, demonstrating that the effect was not driven by movement execution. Consistent with the key-release-locked results, the topographical contrast of beta frequency within the −0.75 to 0.75 s window around keypress onset revealed a frontoparietal pattern (*cluster-based correction, p* = 0.002; Figure S5), confirming the spatial consistency of the effect across temporal alignments. Consistent results were obtained using a conventional frequency-domain metric. Peak beta frequency, computed for the same keypress-aligned window, also differed significantly between long- and short-duration trials (*t_26_*= 2.50, *p* = 0.019; Figure S3B). Together, these results indicate that the beta modulation preceded overt movement and cannot be explained by movement-evoked activity.

**Figure 4.**
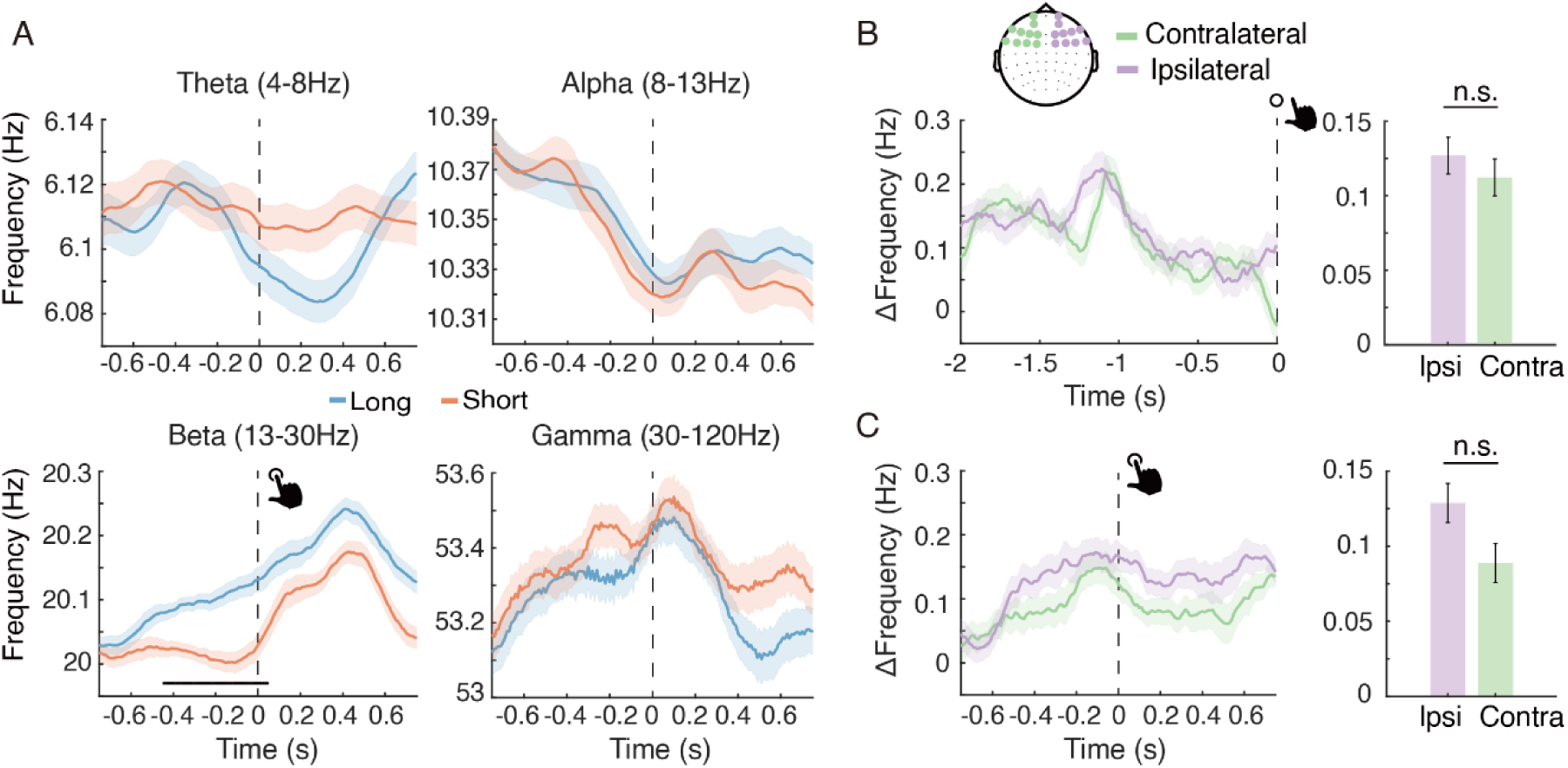
Control analyses excluding movement-evoked and motor-preparatory contributions to beta-frequency modulation. **A**, Instantaneous frequency for the long-and short-duration conditions across four frequency bands: theta (4-8 Hz), alpha (8-13 Hz), beta (13-30 Hz), and gamma (30-120 Hz), time-locked to key press. Significant time points are indicated by the horizontal black bar (*cluster-based correction*, *p* < 0.05). Shaded regions denote ±1 within-subjects SEM. **B,** Comparison of beta-frequency modulation between contralateral and ipsilateral frontoparietal electrode sites relative to the hand used for keypresses in the key-release-locked analysis. The left panel shows the time course of beta-frequency modulation across the pre-release period (−2 to 0 s relative to key release); no significant ipsilateral-contralateral differences were observed (*cluster-based correction,* all *p*s > 0.1). The right panel shows the mean beta-frequency modulation averaged over this period for the two conditions; the ipsilateral and contralateral averages did not differ significantly (*t_26_*= 0.591, *p* = 0.560). **C,** Same as (B), but for the keypress-locked analysis. The left panel shows the time course across the keypress-centered window (−0.75 to 0.75 s relative to keypress onset); again, no significant ipsilateral-contralateral differences were detected (*cluster-based correction,* all *p*s > 0.1). The right panel shows the corresponding mean beta-frequency modulation for the two conditions, which also did not differ significantly (*t_26_*= 1.547, *p* = 0.134).

Although the keypress-locked analysis ruled out the influence of movement-evoked activity, the beta modulation could still, in theory, reflect motor preparation preceding movement execution. To test this possibility, we compared beta-frequency modulation between contralateral and ipsilateral sites within the frontoparietal electrodes, where the main effect was observed, using data analyzed under both key-release- and keypress-locked alignments (Figure 4B and 4C). If the modulation reflected motor preparation, stronger effects would be expected over contralateral motor areas. However, no significant lateralization was detected, indicating that the effect was not effector-specific and therefore unlikely to arise from motor-preparatory processes. As a supplementary validation, we repeated the same contralateral-ipsilateral comparison across all electrodes to ensure that the absence of lateralization was not due to the restricted spatial scope of the analysis (Figure S6). The results were consistent, further confirming that the observed beta modulation was independent of effector-specific motor activity.

Together, these converging findings provide compelling evidence that the observed beta-frequency modulation was independent of motor-related processes, encompassing both movement-evoked and motor-preparatory activity, and instead reflects higher-order timing mechanisms that support self-generated actions.

### Beta-frequency modulation was independent of power and global activation

Next, we examined beta-band power in both key-release- and keypress-locked data to determine whether the frequency modulation was accompanied by changes in oscillatory power. For each alignment, we selected the frontoparietal electrodes that showed significant beta-frequency effects in the corresponding analysis. In both the key-release-locked and keypress-locked analyses, beta power did not differ between the long and short conditions within the time windows where the beta-frequency modulation was significant (−1.95 to −0.94 s relative to key release, and −0.45 to 0.05 s relative to keypress, respectively; Figure 5A and 5B). A beta power difference was observed only in the keypress-locked data between 0.55 and 1.08 s (*cluster-based correction*, *p* = 0.020; Figure 5B), a period outside the frequency effect. This late difference likely reflected the systematic delay in key release between the long- and short-duration conditions rather than a genuine modulation of oscillatory power. To ensure that this dissociation was not caused by spatial restriction, we repeated the analysis across all electrodes. A similar pattern emerged: beta power remained comparable across conditions within the modulation window, and the later difference corresponded to the behavioral timing offset between conditions (Figure S7A). These results indicate that the observed beta-frequency modulation was not driven by either local or global changes in beta power.

**Figure 5.**
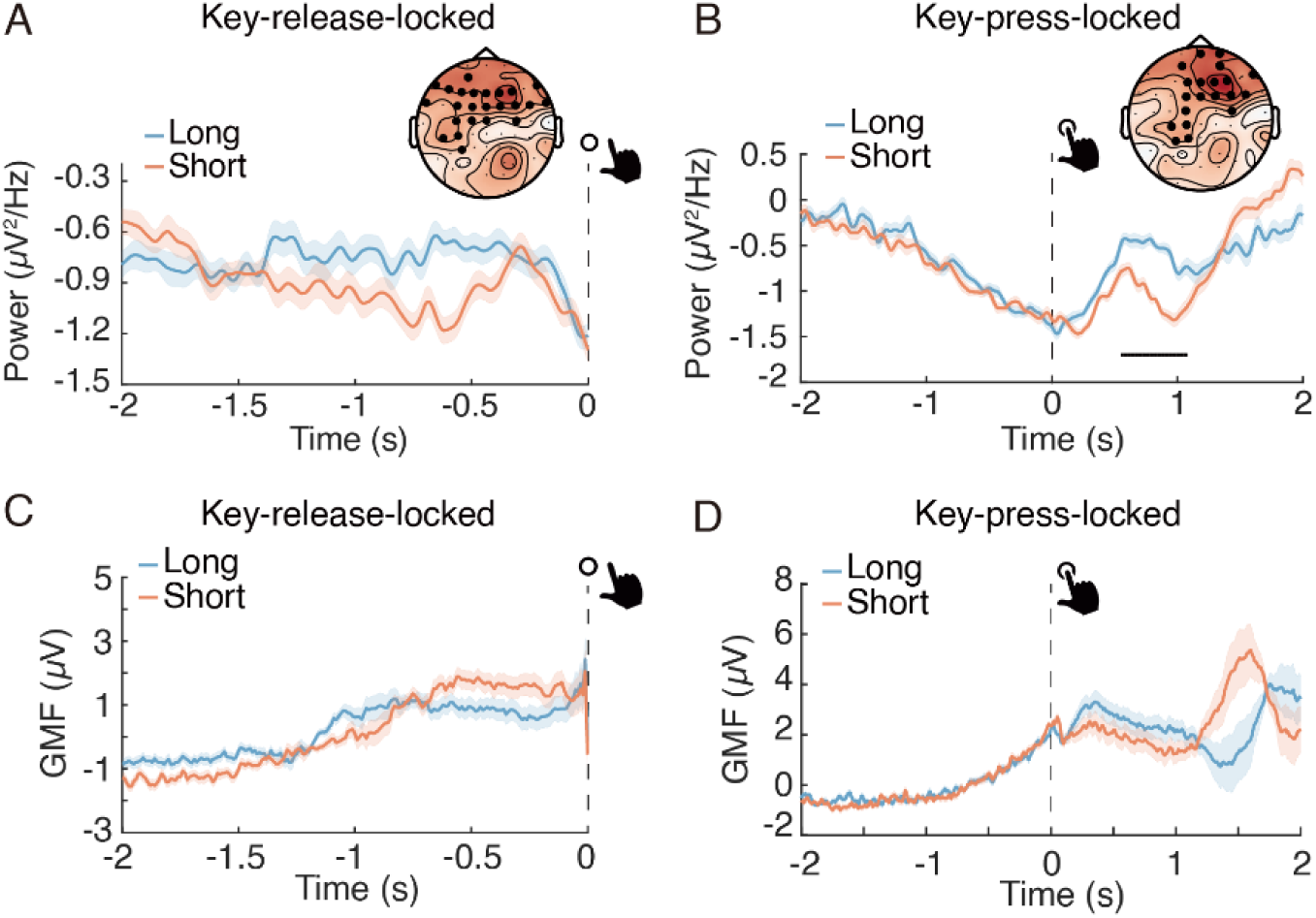
Control analyses excluding power and global activation contributions to beta-frequency modulation. **A,** Beta power for the long- and short-duration conditions averaged over the frontoparietal electrodes (indicated in the top-right scalp map) that showed significant beta-frequency effects in the key-release-locked analysis. **B,** Same as (A), but for the keypress-locked analysis. **C,** Global mean field (GMF) activity for the long-and short-duration conditions in the key-release-locked analysis. **D,** Same as (C), but for the keypress-locked analysis. Significant time points are indicated by horizontal black bars (*cluster-based correction*, *p* < 0.05), and shaded areas represent ±1 within-subject SEM.

We next assessed whether the beta-frequency modulation could be explained by spectral or power-related confounds. After attenuating the 1/f component of the signal using a demodulation procedure and recomputing the instantaneous frequency, the beta-frequency difference between the long and short conditions remained significant (*cluster-based correction*, *p* = 0.047; Figure S7B), confirming that the effect was not due to spectral-slope artifacts.

Finally, we computed the global mean field (GMF) across all electrodes to evaluate potential differences in overall cortical activation. The GMF did not differ between the long and short conditions within the time windows of interest (Figure 5C and 5D). Together, these control analyses demonstrate that the observed beta-frequency modulation cannot be accounted for by differences in beta power, power spectral slope, or global field strength. The findings therefore isolate beta-frequency dynamics as the critical neural signature of self-generated timing.

### Intracranial recordings confirmed the frontoparietal origin of the beta-frequency modulation

To bridge the gap between scalp-level EEG findings and their cortical generators, we next examined intracranial recordings in humans, which provide direct access to neural activity across distributed cortical and subcortical regions with millisecond precision. This approach allowed us to validate the scalp-level beta-frequency modulation and to identify its cortical correlates using invasive recordings. We analyzed sEEG data from two patients with drug-resistant focal epilepsy who were implanted with depth electrodes for clinical monitoring.

Both patients performed the same self-paced time production task as the EEG participants, aiming to produce a 1.5-s target duration. In both cases, self-timed durations were systematically shorter than the target, indicating a consistent underestimation of the 1.5-s duration, similar to that observed in the healthy cohort. For Patient 1, the mean produced duration was 0.83 ± 0.20 s. Trials were divided into tertiles based on the patient’s self-timed durations, yielding short-duration (lower tertile: 0.62 ± 0.06 s) and long-duration (upper tertile: 1.07 ± 0.13 s) conditions. For Patient 2, the mean produced duration was 1.24 ± 0.15 s; the short- and long-duration tertiles averaged 1.08 ± 0.07 s and 1.40 ± 0.10 s, respectively.

At the neural level, we analyzed the instantaneous beta frequency (13-30 Hz) across all implanted contacts for each patient. Averaging across all electrodes provides a comprehensive measure of large-scale beta dynamics, enabling direct comparison with scalp EEG, which similarly reflects activity from distributed cortical sources rather than isolated local generators. Both patients exhibited a pattern consistent with the EEG results: the instantaneous beta frequency was higher during the long-duration than the short-duration condition (Figure 6A and 6C). A cluster-based permutation test revealed temporal clusters distinguishing the two conditions. For Patient 1, a significant cluster was identified from −0.75 to −0.11 s relative to key release (*p* = 0.025; Figure 6A). For Patient 2, a comparable temporal pattern was observed from −1.19 to −0.80 s, approaching significance (*p* = 0.051; Figure 6C).

**Figure 6.**
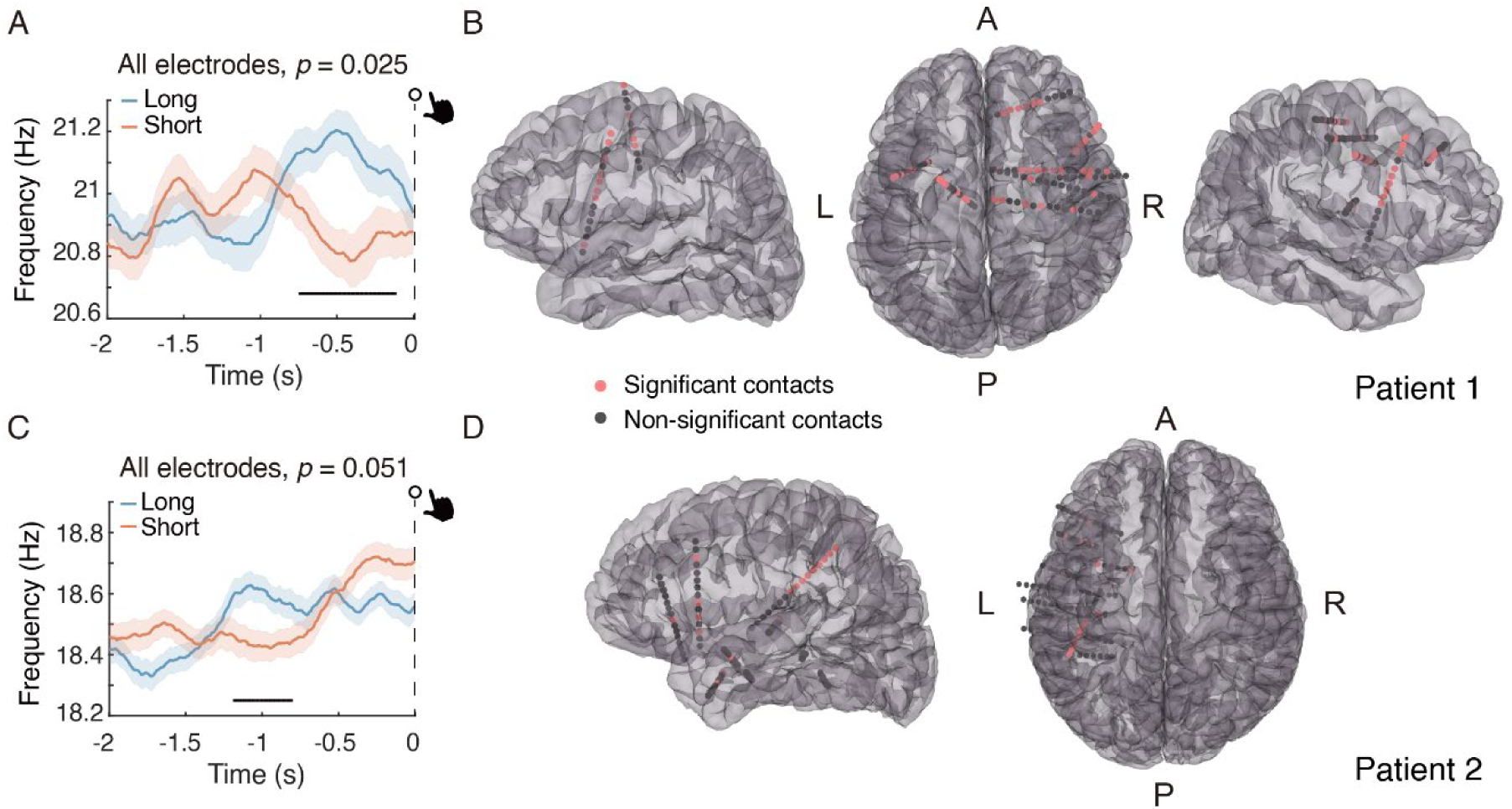
Intracranial recordings of beta-frequency modulation. **A,** Instantaneous beta frequency (13-30 Hz) averaged across all implanted electrode contacts for Patient 1 in the long- and short-duration conditions. The horizontal black bar indicates time points showing a significant difference between conditions (*cluster-based correction*, *p* = 0.025). **B,** Spatial distribution of electrode contacts for Patient 1. Colored contacts mark electrode contacts with significantly higher beta frequency (long > short) during the clusters shown in (A), and gray contacts represent non-significant contacts. **C,** Same as (A), but for Patient 2. No cluster survived correction for multiple comparisons, although a trend-level difference was observed (*cluster-based correction*, *p* = 0.051). **D,** Same as (B), but for Patient 2, with colored contacts defined based on the clusters shown in (C). Shaded areas represent ±1 within-subject SEM. Brain renderings correspond to each patient’s native anatomy and are shown for visualization only. Anatomical orientation markers are indicated as A (anterior), P (posterior), L (left), and R (right).

We then localized the electrode contacts that exhibited significantly higher beta frequency in the long-duration condition within the time windows of interest, defined by the significant or near-significant clusters identified above. The significant contacts were primarily located in the frontal and parietal cortices (Figure 6B and 6D), closely matching the frontoparietal distribution of the beta-frequency modulation observed in the scalp EEG data.

Together, these intracranial findings confirm that the beta-frequency modulation observed at the scalp originates from distributed frontoparietal cortical networks, providing convergent invasive evidence that this oscillatory mechanism supports the generation of self-timed actions.

## Discussion

We identify a neural mechanism supporting self-generated timing based on moment-to-moment modulation of instantaneous beta frequency. Higher beta frequency (13-30 Hz) reliably predicted longer produced durations, both within individuals on a trial-by-trial basis and across individuals at a trait level, and was localized to a distributed frontoparietal network. This relationship was robust to multiple motor- and signal-related confounds and was confirmed by convergent intracranial recordings. Together, these findings establish instantaneous beta frequency as a scalable neural correlate of self-generated timing.

A central implication of this result concerns the nature of internal timing mechanisms. Classical pacemaker-accumulator models predict that faster oscillatory rhythms should lead to shorter subjective durations through increased temporal sampling(Buhusi & Meck, 2005; Gibbon, 1977; Treisman, 1963). In contrast, our data reveal the opposite pattern: higher instantaneous beta frequency (IBF) is associated with longer self-generated time durations. This dissociation indicates that beta frequency does not conform to the core predictions of internal clock models, but instead reflects a distinct control process that biases the persistence and termination of ongoing actions, favoring a stabilizing control account over pacemaker-accumulator explanations.

This interpretation aligns with the “status quo” hypothesis of beta oscillations, which proposes that beta activity supports the maintenance of existing cognitive or motor states and resists change(Engel & Fries, 2010; Kilavik et al., 2013). Empirical work supports this view across multiple domains, demonstrating that increased beta-band synchrony preferentially reinforces existing motor states while impairing the initiation and execution of new movements(Gilbertson et al., 2005; Pogosyan et al., 2009), that beta activity indexes the stabilization and endogenous reactivation of internal sensorimotor goals and task-relevant representations(Spitzer & Haegens, 2017; Tan et al., 2016), and that beta-band oscillations mediate rhythmic top-down control from prefrontal regions to motor cortex during action suppression and response inhibition(Picazio et al., 2014; Stolk et al., 2019; Swann et al., 2013). In the present task, instantaneous beta frequency is specifically linked to the timing of voluntary key release. Higher beta frequency may increase the frequency of stabilizing control updates within a given time interval, thereby sustaining the ongoing action tendency for longer before transition. As a result, elevated beta frequency is associated with delayed release times and longer produced durations. These findings suggest that beta-frequency dynamics regulate when an internally generated action is terminated during self-paced timing by modulating the density of intrinsic control updates.

Importantly, this control-based account is fully compatible with state-dependent and population-based models of timing, which conceptualize time as emerging from the evolution of neural network states rather than from explicit clock-like mechanisms(Buonomano & Laje, 2010; Buonomano & Maass, 2009; Paton & Buonomano, 2018). Within these frameworks, temporal information is carried by the continuous dynamics of population activity, rather than by the accumulation of discrete temporal units. From this perspective, instantaneous beta frequency need not encode elapsed time itself, but may instead index the degree to which ongoing network states are stabilized during duration generation. In doing so, beta-frequency dynamics provide a control signal that regulates the stability of internal state trajectories, complementing state-dependent representations of elapsed time and linking oscillatory control processes to population-based timing mechanisms.

The specificity of our results to the beta band further supports the view that beta-frequency dynamics play a privileged role in the regulation of self-generated timing. Despite using identical analytic pipelines, we observed no reliable modulations of instantaneous frequency in the theta, alpha, or gamma bands across conditions. This dissociation suggests that the observed timing-related effects are not a general property of oscillatory activity or signal variability, but are functionally tied to the beta rhythm. While alpha and theta oscillations have been implicated in attentional sampling, working memory, temporal expectation under externally cued conditions(Cecere et al., 2015; Han, Zhang, et al., 2023; Samaha & Postle, 2015; Shen et al., 2019; VanRullen, 2016; Wolinski et al., 2018; Xu et al., 2025) and time perception(Anliker, 1963; Frisoni et al., 2025; Grabot et al., 2019; Legg, 1968; Mioni et al., 2020; Treisman, 1984; Wutz, 2024), the present task required purely endogenous control of time, under which beta-band dynamics appear uniquely suited to sustain internal timing goals.

A striking feature of our findings is their consistency across individuals, analytic approaches and recording modalities. Participants who exhibited larger beta-frequency differences between long- and short-duration trials also showed greater behavioral timing shifts, indicating that beta-frequency modulation captures a stable, trait-like component of individual timing dynamics. Importantly, these trait-level relationships arise from systematic aggregation of moment-to-moment beta-frequency fluctuations rather than reflecting a static individual parameter. Consistent effects were also observed using an FFT-based estimate of peak beta frequency, suggesting that the beta-duration relationship generalizes across analytic approaches. Convergent intracranial recordings further confirmed that this relationship originates from distributed cortical networks, providing direct evidence that the beta-frequency modulation observed at the scalp reflects genuine cortical dynamics rather than measurement artifacts(Han, Shen, et al., 2023; Parvizi & Kastner, 2018). Together, this cross-participant and cross-modality convergence underscores the generality and robustness of beta-frequency dynamics as a neural mechanism supporting self-generated timing, supported by analytical approaches that resolve fine-grained frequency fluctuations beyond traditional power- or phase-based measures.

Several limitations should be acknowledged. While our findings support a control-based account of beta-frequency dynamics in self-generated timing, the present results are correlational in nature. Future studies combining frequency-specific neuromodulation with closed-loop designs will be essential to establish a causal role for instantaneous beta frequency in shaping subjective time. In addition, whether similar beta-frequency dynamics generalize to externally cued or more complex temporal contexts remains an important question for future work.

Together, our findings indicate that instantaneous beta frequency governs self-generated time control—the regulation of when an internally guided action should persist or terminate. Rather than functioning as a neural clock, higher beta frequency provides more frequent stabilizing control within a given time interval, delaying action termination and prolonging produced durations. In this framework, internally generated timing is shaped not by the accumulation of temporal ticks, but by the density of intrinsic control updates that regulate action persistence.

## Methods

### Participants

Twenty-eight healthy, right-handed adults (17 females; mean age = 20.4 ± 1.9 years) were enrolled in the scalp EEG experiment. One participant was excluded prior to analysis due to an insufficient number of artifact-free trials, yielding a final sample of 27. The target sample size was determined a priori to ensure ≥80 % statistical power to detect moderate-to-large effect sizes (Cohen’s *d* > 0.6). In parallel, three right-handed patients with drug-resistant focal epilepsy (one female; mean age = 28 ± 1.4 years) undergoing invasive monitoring participated in the study. One patient was recruited from the Guangdong Second Provincial General Hospital and two patients were recruited from the Second Affiliated Hospital of Guangzhou Medical University. Depth electrodes were implanted solely for clinical seizure localization; no additional electrodes were placed for research purposes. One patient’s data was excluded from further analysis because of excessive epileptiform activity, resulting in a final intracranial sample of two patients. Following implantation, patients remained under continuous video-EEG surveillance in the hospital for approximately two weeks and completed the identical experimental task used in the healthy cohort. All participants reported normal or corrected-to-normal vision and no history of psychiatric or neurological disorders (aside from epilepsy in the patient group). Written informed consent was obtained from every individual in accordance with the Declaration of Helsinki. The study protocol was reviewed and approved by the Ethics Committee of the School of Psychology, South China Normal University.

### Experimental Design

Participants performed a self-paced time-production task while fixating a centrally presented green dot on a uniform gray background, with continuous EEG recorded throughout the task. Each trial commenced after a mandatory 3-s intertrial interval. Participants initiated a trial by pressing a thin-film pressure sensor with their right index finger; if this occurred before the 3-s interval had elapsed, an auditory warning (“pressing key too soon”) sounded and the trial automatically restarted.

Upon valid initiation, participants estimated a 1.5 s duration by continuing to hold down the sensor and then released it to terminate the trial. They were instructed to aim for as precise timing as possible, without reference to any external time standard and while refraining from overt strategies such as counting or foot tapping. Auditory feedback was provided only for significant timing deviations: a beep signaled productions shorter than 0.5 s or longer than 3 s, whereas releases within the 0.5-3 s window produced no sound. Participants were not informed about the specific feedback window; they were only told that auditory feedback would be provided for large timing errors. To minimize device latency and ensure accurate timestamps, the pressure sensor was interfaced with an Arduino Uno microcontroller, and all press/release events were recorded using Presentation software (Neurobehavioral Systems; http://www.neurobs.com).

Prior to the main session, participants completed a practice block to familiarize themselves with the task structure and the two feedback sounds. The main experiment comprised 400 trials per participant, with brief rest breaks every 50 trials.

### Recording and Preprocessing of EEG Data

Continuous EEG was acquired at 1,024 Hz using a NeuronScan SynAmps2 system with a 64 - channel Quick-Cap fitted with Ag/AgCl electrodes. Four additional electrodes recorded horizontal and vertical electrooculograms (EOG) around the eyes. Electrode impedances were kept below 5 kΩ, and all channels were referenced online to FCz.

Offline preprocessing and analysis were performed in MATLAB using the FieldTrip toolbox(Oostenveld et al., 2011) (www.fieldtriptoolbox.org) alongside custom scripts. The raw data were downsampled to 512 Hz, notch-filtered at 50 Hz to remove power-line noise, and then re-referenced to the average of all scalp electrodes. Data were epoched from −3,000 ms to +3,000 ms relative to keypress onset and baseline-corrected by subtracting the mean amplitude during the −2,000 to −1,000 ms pre-keypress window. Trials with premature keypresses or beep - triggered errors—and the immediately following trial—were excluded. Productions exceeding three standard deviations from each participant’s mean duration were then removed. Remaining epochs underwent visual inspection to eliminate ocular and muscle artifacts. Trials shorter than 750 ms were discarded to ensure a sufficient post-response analysis window. The durations of all rejected trials (across all criteria) were 1.196 ± 0.470 s (mean ± SD).

### Recording and Preprocessing of Intracranial Data

Depending on clinical requirements, semi-rigid platinum-iridium depth electrodes (0.8 mm diameter, 2 mm length, 1.5 mm inter-contact spacing) with 7-19 contacts each were stereotactically implanted. After implantation, patients remained under continuous video-EEG monitoring in their hospital rooms for approximately two weeks. SEEG signals were acquired using a 256-channel Nihon Kohden Neurofax 1200A digital system in bipolar montage at a sampling rate of 2,000 Hz.

Electrode localization followed established procedures(Yang et al., 2024). Post-implant CT scans were coregistered with pre-implant T1-weighted MRI using the FMRIB Software Library(Jenkinson et al., 2012). Contacts were identified and reconstructed with the sEEG Assistant(Narizzano et al., 2017) implemented in 3D Slicer(Fedorov et al., 2012), and electrode positions were visualized in 3D(Fischl, 2012).

Preprocessing steps paralleled those used for the scalp EEG dataset. Raw recordings were downsampled to 1,000 Hz, notch-filtered at 50 Hz to attenuate line noise, and re-referenced to the common average. Data were epoched from −3,000 to +3,000 ms relative to keypress onset and baseline-corrected using the −2,000 to −1,000 ms window. For analyses aligned to key release, epochs spanned −2,000 to 0 ms. Trials contaminated by epileptiform activity were identified through visual inspection and excluded.

### Behavioral and Neural Data Analysis

For each participant, self-timed durations (key-release times) were sorted and divided into tertiles. Trials falling below the first tertile were classified as short-duration trials, whereas those exceeding the third tertile were classified as long-duration trials. Mean self-timed durations (± SD) were calculated for each participant separately for the short- and long-duration conditions. Paired-samples t-tests were conducted to compare conditions at the group level. To characterize individual variability, we further examined the distribution of self-timed durations across participants, highlighting differences between subjective timing and the objective 1.5-s target duration.

Instantaneous frequency and power analyses followed established procedures(Cohen, 2014b). Specifically, instantaneous-frequency analyses were performed using data from - 2,000 to 0 ms relative to key-release and from −750 to 750 ms relative to key-press onset, thereby avoiding contamination from release-related activity. Each epoch was band-pass filtered between 13-30 Hz (beta band) or, in control analyses, within 4-8 Hz (theta), 8-13 Hz (alpha), or 30-120 Hz (gamma) using a zero-phase, plateau-shaped finite-impulse-response filter with a 15% transition width. A Hilbert transform was then applied to extract the phase-angle and amplitude time series on each trial. The temporal derivative of the instantaneous Hilbert phase (scaled by sampling rate / 2π) yielded instantaneous frequency in hertz, whereas the squared amplitude reflected oscillatory power. To suppress non-physiological spikes induced by noise in the phase derivative, the instantaneous-frequency estimates were median-filtered ten times using windows ranging from 10 to 400 ms, following prior recommendations(Cohen, 2014b; Samaha & Postle, 2015).

As an additional frequency-domain analysis, peak beta frequency (PBF) was estimated from scalp EEG data using standard spectral analysis procedures implemented in the FieldTrip toolbox. Power spectra were computed over the 2-50 Hz frequency range using a Hanning taper, and zero-padding was applied to extend the data length to 10 s in order to increase frequency sampling density. Power spectra were averaged across all scalp electrodes to obtain a global spectral profile for each condition. PBF was defined as the frequency corresponding to the maximum power within the beta band (13-30 Hz) and was subsequently compared between short- and long-duration conditions.

To control for potential influences of the aperiodic 1/f component, we used the FOOOF algorithm(Donoghue et al., 2020) in the FieldTrip toolbox and reconstructed demodulated time-domain signals with attenuated 1/f structure(Samaha & Cohen, 2022). To minimize edge artifacts from filtering, each trial was mirrored in time and appended to its original segment before Fourier transformation. The amplitude and phase spectra were extracted via FFT; the FOOOF model fit and removed the aperiodic component from the power spectrum, producing a flattened spectrum. The corrected amplitude spectrum was recombined with the original phase spectrum to generate complex values, which were then inverse-transformed to yield the demodulated signal. Instantaneous-frequency analysis was subsequently repeated on these demodulated signals within the fixed 13-30 Hz range for all participants.

Time-frequency power was estimated using a short-time Fourier transform with a single Hanning taper, implemented in the FieldTrip toolbox. Power spectra were computed on all trials. Frequencies of interest ranged from 2 to 50 Hz in 1-Hz steps, and the number of cycles used for analysis increased linearly from 2 to 7 across the frequency range to balance temporal and spectral resolution. Analyses were performed separately for key-release-locked (−2,000 to 0 ms) and key-press-locked (−2,000 to 2,000 ms) epochs. For each condition, beta-band (13-30 Hz) power was averaged across frequencies within this range to obtain a single power estimate per time point.

To quantify global cortical activity without assuming any specific source configuration, the global mean field (GMF) was also computed using the FieldTrip toolbox. GMF was calculated as the spatial standard deviation of scalp potentials across all electrodes at each time point, providing a reference-free measure of overall neural synchrony. GMF time series were derived for both key-release-locked and key-press-locked epochs using the same preprocessing pipeline described above. For each participant, GMF traces were averaged within the short- and long-duration conditions to obtain condition-specific mean waveforms.

### Statistical Analysis

Within-subject comparisons between the short- and long-duration conditions were conducted using two-tailed paired-samples t-tests for scalp EEG data and one-tailed independent-samples t-tests for intracranial (sEEG) data, reflecting the directional hypothesis derived from group-level effects. To assess inter-individual relationships, Pearson correlations were computed between EEG-derived neural metrics (e.g., instantaneous beta frequency, IBF) and behavioral measures such as self-timed duration.

Temporal-level statistical evaluation of instantaneous frequency and other neural measures was performed using a nonparametric cluster-based permutation test(Maris & Oostenveld, 2007), which effectively controls for multiple comparisons across time and sensors. For the group-level EEG analysis, data from all participants were pooled, whereas sEEG data were analyzed at the single-subject level. At each time point and electrode, t-values were computed for the contrast between conditions, and samples exceeding a two-sided threshold of p < 0.05 were provisionally marked. Adjacent time points meeting this criterion were grouped into temporal clusters for the time-course analyses, whereas neighboring electrodes were grouped into spatial clusters for the topographical analyses. In each case, the sum of t-values within a cluster was used as the cluster-level statistic.

To obtain the null distribution, condition labels were randomly shuffled 1,000 times, and the maximum cluster-level statistic from each permutation was retained. An observed cluster was deemed significant if its cluster-level statistic exceeded the 95th percentile of the permutation distribution (α = 0.05). This procedure was applied consistently across all time-resolved EEG and sEEG analyses reported in the study.

## Data and code availability

The data reported in this study and the custom analysis code have been deposited at the Open Science Framework: https://osf.io/r4dz9/. They have been made available for reuse or distribution in the public domain, as per the criteria set forth by the institute and in compliance with the ethical committee’s endorsement.

## Acknowledgments

This work was supported by the National Key Research and Development Program of China (2025YFE0213500); the National Natural Science Foundation of China (32471144); the Research Center for Brain Cognition and Human Development, Guangdong, China (2024B0303390003); the MOE Project of Key Research Institute of Humanities and Social Sciences in Universities (22JJD190006); and the Striving for the First-Class, Improving Weak Links and Highlighting Features (SIH) Key Discipline Program for Psychology at South China Normal University. We thank Dr. Vincenzo Romei for helpful discussions and insightful comments on the manuscript.

## Reference

Anliker, J. (1963). Variations in Alpha Voltage of the Electroencephalogram and Time Perception. Science, 140(3573), 1307–1309. 10.1126/science.140.3573.1307

Bartolo, R. on, Prado, L., & Merchant, H. (2014). Information Processing in the Primate Basal Ganglia during Sensory-Guided and Internally Driven Rhythmic Tapping. The Journal of Neuroscience, 34(11), 3910–3923. 10.1523/JNEUROSCI.2679-13.2014

Buhusi, C. V., & Meck, W. H. (2005). What makes us tick? Functional and neural mechanisms of interval timing. Nature Reviews Neuroscience, 6(10), 755–765. 10.1038/nrn1764

Buonomano, D. V., & Laje, R. (2010). Population clocks: Motor timing with neural dynamics. Trends in Cognitive Sciences, 14(12), 520–527. 10.1016/j.tics.2010.09.002

Buonomano, D. V., & Maass, W. (2009). State-dependent computations: Spatiotemporal processing in cortical networks. Nature Reviews Neuroscience, 10(2), 113–125. 10.1038/nrn2558

Cecere, R., Rees, G., & Romei, V. (2015). Individual differences in alpha frequency drive crossmodal illusory perception. Current Biology, 25(2), 231–235. 10.1016/j.cub.2014.11.034

Cohen, M. X. (2014a). Analyzing Neural Time Series Data: Theory and Practice. MIT Press.

Cohen, M. X. (2014b). Fluctuations in oscillation frequency control spike timing and coordinate neural networks. Journal of Neuroscience, 34(27), 8988–8998. 10.1523/JNEUROSCI.0261-14.2014

Donoghue, T., Haller, M., Peterson, E. J., Varma, P., Sebastian, P., Gao, R., Noto, T., Lara, A. H., Wallis, J. D., Knight, R. T., Shestyuk, A., & Voytek, B. (2020). Parameterizing neural power spectra into periodic and aperiodic components. Nature Neuroscience, 23(12), 1655–1665. 10.1038/s41593-020-00744-x

Engel, A. K., & Fries, P. (2010). Beta-band oscillations—Signalling the status quo? Current Opinion in Neurobiology, 20(2), 156–165. 10.1016/j.conb.2010.02.015

Fedorov, A., Beichel, R., Kalpathy-Cramer, J., Finet, J., Fillion-Robin, J.-C., Pujol, S., Bauer, C., Jennings, D., Fennessy, F., Sonka, M., Buatti, J., Aylward, S., Miller, J. V., Pieper, S., & Kikinis, R. (2012). 3D Slicer as an image computing platform for the Quantitative Imaging Network. Magnetic Resonance Imaging, Quantitative Imaging in Cancer, 30(9), 1323–1341. 10.1016/j.mri.2012.05.001

Feurra, M., Bianco, G., Santarnecchi, E., Del Testa, M., Rossi, A., & Rossi, S. (2011). Frequency-Dependent Tuning of the Human Motor System Induced by Transcranial Oscillatory Potentials. The Journal of Neuroscience, 31(34), 12165–12170. 10.1523/JNEUROSCI.0978-11.2011

Fischl, B. (2012). FreeSurfer. NeuroImage, 20 YEARS OF fMRI, 62(2), 774–781. 10.1016/j.neuroimage.2012.01.021

Frisoni, M., Tarasi, L., Borgomaneri, S., & Romei, V. (2025). The relationship between individual alpha frequency and time perception: Testing the internal clock versus the sampling rate hypothesis. Cortex, 192, 183–195. 10.1016/j.cortex.2025.09.008

Fujioka, T., Trainor, L. J., Large, E. W., & Ross, B. (2012). Internalized Timing of Isochronous Sounds Is Represented in Neuromagnetic Beta Oscillations. The Journal of Neuroscience, 32(5), 1791–1802. 10.1523/JNEUROSCI.4107-11.2012

Gibbon, J. (1977). Scalar expectancy theory and Weber’s law in animal timing. Psychological Review, 84(3), 279–325. 10.1037/0033-295X.84.3.279

Gilbertson, T., Lalo, E., Doyle, L., Lazzaro, V. D., Cioni, B., & Brown, P. (2005). Existing Motor State Is Favored at the Expense of New Movement during 13-35 Hz Oscillatory Synchrony in the Human Corticospinal System. Journal of Neuroscience, 25(34), 7771–7779. 10.1523/JNEUROSCI.1762-05.2005

Grabot, L., Kononowicz, T. W., Tour, T. D. e, Gramfort, A., Doy\‘ ere, V. erie, & Wassenhove, V. (2019). The Strength of Alpha–Beta Oscillatory Coupling Predicts Motor Timing Precision. Journal of Neuroscience, 39(17), 3277–3291. 10.1523/JNEUROSCI.2473-18.2018

Han, B., Shen, L., & Chen, Q. (2023). How Can I Combine Data from fMRI, EEG, and Intracranial EEG? In N. Axmacher (Ed.), Intracranial EEG: A Guide for Cognitive Neuroscientists (pp. 239–256). Springer International Publishing. 10.1007/978-3-031-20910-9_15

Han, B., Zhang, Y., Shen, L., Mo, L., & Chen, Q. (2023). Task demands modulate pre-stimulus alpha frequency and sensory template during bistable apparent motion perception. Cerebral Cortex, 33(5), 1679–1692. 10.1093/cercor/bhac165

Jenkinson, M., Beckmann, C. F., Behrens, T. E. J., Woolrich, M. W., & Smith, S. M. (2012). FSL. NeuroImage, 62(2), 782–790. 10.1016/j.neuroimage.2011.09.015

Kilavik, B. E., Zaepffel, M., Brovelli, A., MacKay, W. A., & Riehle, A. (2013). The ups and downs of beta oscillations in sensorimotor cortex. Experimental Neurology, 245, 15–26. 10.1016/j.expneurol.2012.09.014

Kononowicz, T. W., & van Rijn, H. (2015). Single trial beta oscillations index time estimation. Neuropsychologia, 75, 381–389. 10.1016/j.neuropsychologia.2015.06.014

Kononowicz, T. W., van Rijn, H., & Meck, W. H. (2018). Timing and Time Perception: A Critical Review of Neural Timing Signatures Before, During, and After the To-Be-Timed Interval. In J. T. Wixted (Ed.), Stevens’ Handbook of Experimental Psychology and Cognitive Neuroscience (pp. 1–38). John Wiley & Sons, Inc. 10.1002/9781119170174.epcn114

Kulashekhar, S., Pekkola, J., Palva, J. M., & Palva, S. (2016). The role of cortical beta oscillations in time estimation. Human Brain Mapping, 37(9), 3262–3281. 10.1002/hbm.23239

Legg, C. F. (1968). Alpha rhythm and time judgments. Journal of Experimental Psychology, 78(1), 46–49. 10.1037/h0026149

Little, S., Bonaiuto, J., Barnes, G., & Bestmann, S. (2019). Human motor cortical beta bursts relate to movement planning and response errors. PLOS Biology, 17(10), e3000479. 10.1371/journal.pbio.3000479

Maris, E., & Oostenveld, R. (2007). Nonparametric statistical testing of EEG- and MEG-data. Journal of Neuroscience Methods, 164(1), 177–190. 10.1016/j.jneumeth.2007.03.024

Merchant, H., Harrington, D. L., & Meck, W. H. (2013). Neural Basis of the Perception and Estimation of Time. Annual Review of Neuroscience, 36(Volume 36, 2013), 313–336. 10.1146/annurev-neuro-062012-170349

Milton, A., & Pleydell-Pearce, C. W. (2016). The phase of pre-stimulus alpha oscillations influences the visual perception of stimulus timing. Neuroimage, 133, 53–61. 10.1016/j.neuroimage.2016.02.065

Mioni, G., Shelp, A., Stanfield-Wiswell, C. T., Gladhill, K. A., Bader, F., & Wiener, M. (2020). Modulation of Individual Alpha Frequency with tACS shifts Time Perception. Cerebral Cortex Communications, 1(1), tgaa064. 10.1093/texcom/tgaa064

Morrow, A., Wilson, M., Geller-Montague, M., Soldano, S., Hajidamji, S., & Samaha, J. (2024). Individual Alpha Frequency Predicts the Sensitivity of Time Perception. 10.1101/2024.12.16.628734

Narizzano, M., Arnulfo, G., Ricci, S., Toselli, B., Tisdall, M., Canessa, A., Fato, M. M., & Cardinale, F. (2017). SEEG assistant: A 3DSlicer extension to support epilepsy surgery. BMC Bioinformatics, 18(1), 124. 10.1186/s12859-017-1545-8

Oostenveld, R., Fries, P., Maris, E., & Schoffelen, J.-M. (2011). FieldTrip: Open source software for advanced analysis of MEG, EEG, and invasive electrophysiological data. Computational Intelligence and Neuroscience, 2011, 156869. 10.1155/2011/156869

Parvizi, J., & Kastner, S. (2018). Promises and limitations of human intracranial electroencephalography. Nature Neuroscience, 21(4), 474–483. 10.1038/s41593-018-0108-2

Paton, J. J., & Buonomano, D. V. (2018). The Neural Basis of Timing: Distributed Mechanisms for Diverse Functions. Neuron, 98(4), 687–705. 10.1016/j.neuron.2018.03.045

Picazio, S., Veniero, D., Ponzo, V., Caltagirone, C., Gross, J., Thut, G., & Koch, G. (2014). Prefrontal Control over Motor Cortex Cycles at Beta Frequency during Movement Inhibition. Current Biology, 24(24), 2940–2945. 10.1016/j.cub.2014.10.043

Pogosyan, A., Gaynor, L. D., Eusebio, A., & Brown, P. (2009). Boosting Cortical Activity at Beta-Band Frequencies Slows Movement in Humans. Current Biology, 19(19), 1637–1641. 10.1016/j.cub.2009.07.074

Samaha, J., & Cohen, M. X. (2022). Power spectrum slope confounds estimation of instantaneous oscillatory frequency. NeuroImage, 250, 118929. 10.1016/j.neuroimage.2022.118929

Samaha, J., & Postle, B. R. (2015). The Speed of Alpha-Band Oscillations Predicts the Temporal Resolution of Visual Perception. Current Biology, 25(22), 2985–2990. 10.1016/j.cub.2015.10.007

Schilberg, L., Engelen, T. ee, ten Oever, S., Schuhmann, T., de Gelder, B., de Graaf, T. A., & Sack, A. T. (2018). Phase of beta-frequency tACS over primary motor cortex modulates corticospinal excitability. Cortex, 103, 142–152. 10.1016/j.cortex.2018.03.001

Shen, L., Han, B., Chen, L., & Chen, Q. (2019). Perceptual inference employs intrinsic alpha frequency to resolve perceptual ambiguity. PLoS Biology, 17(3), e3000025.

Sherman MA, Lee S, Law R, Haegens S, Thorn CA, Hämäläinen MS, Moore CI, Jones SR. (2016). Neural mechanisms of transient neocortical beta rhythms: Converging evidence from humans, computational modeling, monkeys, and mice. Proceedings of the National Academy of Sciences, 113(33). 10.1073/pnas.1604135113

Simen, P., Balci, F., deSouza, L., Cohen, J. D., & Holmes, P. (2011). A Model of Interval Timing by Neural Integration. Journal of Neuroscience, 31(25), 9238–9253. 10.1523/JNEUROSCI.3121-10.2011

Spitzer, B., & Haegens, S. (2017). Beyond the Status Quo: A Role for Beta Oscillations in Endogenous Content (Re)Activation. eNeuro, 4(4), ENEURO.0170-17.2017. 10.1523/ENEURO.0170-17.2017

Stolk, A., Brinkman, L., Vansteensel, M. J., Aarnoutse, E., Leijten, F. S., Dijkerman, C. H., Knight, R. T., de Lange, F. P., & Toni, I. (2019). Electrocorticographic dissociation of alpha and beta rhythmic activity in the human sensorimotor system. eLife, 8, e48065. 10.7554/eLife.48065

Swann, N. C., Tandon, N., Pieters, T. A., & Aron, A. R. (2013). Intracranial electroencephalography reveals different temporal profiles for dorsal- and ventro-lateral prefrontal cortex in preparing to stop action. Cerebral Cortex, 23(10), 2479–2488. 10.1093/cercor/bhs245

Tan, H., Wade, C., & Brown, P. (2016). Post-Movement Beta Activity in Sensorimotor Cortex Indexes Confidence in the Estimations from Internal Models. The Journal of Neuroscience, 36(5), 1516–1528. 10.1523/JNEUROSCI.3204-15.2016

Treisman, M. (1963). Temporal discrimination and the indifference interval: Implications for a model of the “internal clock”. Psychological Monographs: General and Applied, 77(13), 1–31. 10.1037/h0093864

Treisman, M. (1984). Temporal Rhythms and Cerebral Rhythms. Annals of the New York Academy of Sciences, 423(1 Timing and Ti), 542–565. 10.1111/j.1749-6632.1984.tb23458.x

Tsao, A., Yousefzadeh, S. A., Meck, W. H., Moser, M.-B., & Moser, E. I. (2022). The neural bases for timing of durations. Nature Reviews Neuroscience, 23(11), 646–665. 10.1038/s41583-022-00623-3

VanRullen, R. (2016). Perceptual Cycles. Trends in Cognitive Sciences, 20(10), 723–735. 10.1016/j.tics.2016.07.006

Wiener, M., Parikh, A., Krakow, A., & Coslett, H. B. (2018). An Intrinsic Role of Beta Oscillations in Memory for Time Estimation. Scientific Reports, 8(1), 7992. 10.1038/s41598-018-26385-6

Wolinski, N., Cooper, N. R., Sauseng, P., & Romei, V. (2018). The speed of parietal theta frequency drives visuospatial working memory capacity. PLOS Biology, 16(3), e2005348. 10.1371/journal.pbio.2005348

Wutz, A. (2024). Alpha Oscillations Create the Illusion of Time. Journal of Cognitive Neuroscience, 1–9. 10.1162/jocn_a_02029

Xu, M., Han, B., Chen, Q., & Shen, L. (2025). Auditory stimuli extend the temporal window of visual integration by modulating alpha-band oscillations. eLife. http://biorxiv.org/lookup/doi/10.1101/2024.01.31.578121

Yang, J., Shen, L., Long, Q., Li, W., Zhang, W., Chen, Q., & Han, B. (2024). Electrical stimulation induced self-related auditory hallucinations correlate with oscillatory power change in the default mode network. Cerebral Cortex, 34(1), bhad473. 10.1093/cercor/bhad473

