## Supplementary material for "Instantaneous Beta Frequency Regulates Self-Generated Timing in Humans": Figure S1

**Supplementary Files**

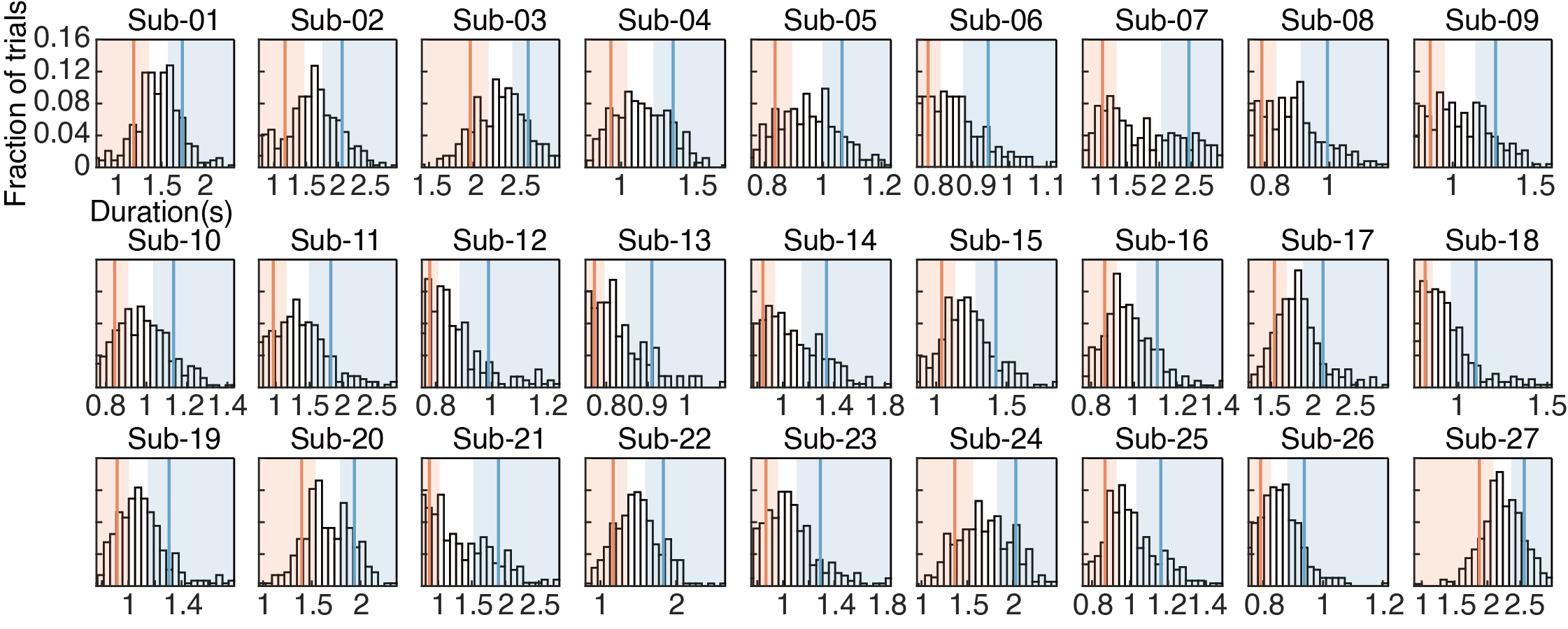

**Figure S1. Distribution of produced durations across participants.** Red and blue rectangles indicate the ranges of short and long trials selected for analysis, respectively, and the corresponding red and blue vertical lines denote their mean values.

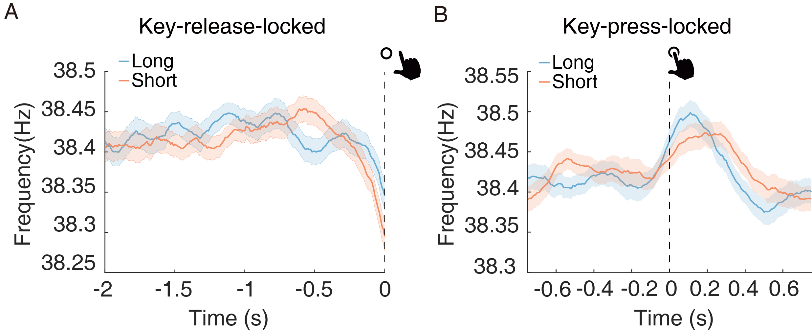

**Figure S2.** **Relationship between instantaneous frequency and self-timed duration in the low-gamma band (30-50 Hz).** **A**, Instantaneous frequency for the long- and short-duration conditions in the key-release-locked analysis. No significant differences were observed between conditions (*cluster-based correction*, all *p*s > 0.1). **B,** Same as (A), but for the keypress-locked analysis. Again, no significant differences were detected between conditions (*cluster-based correction*, all *p*s > 0.1). Shaded regions denote ±1 within-subjects SEM.

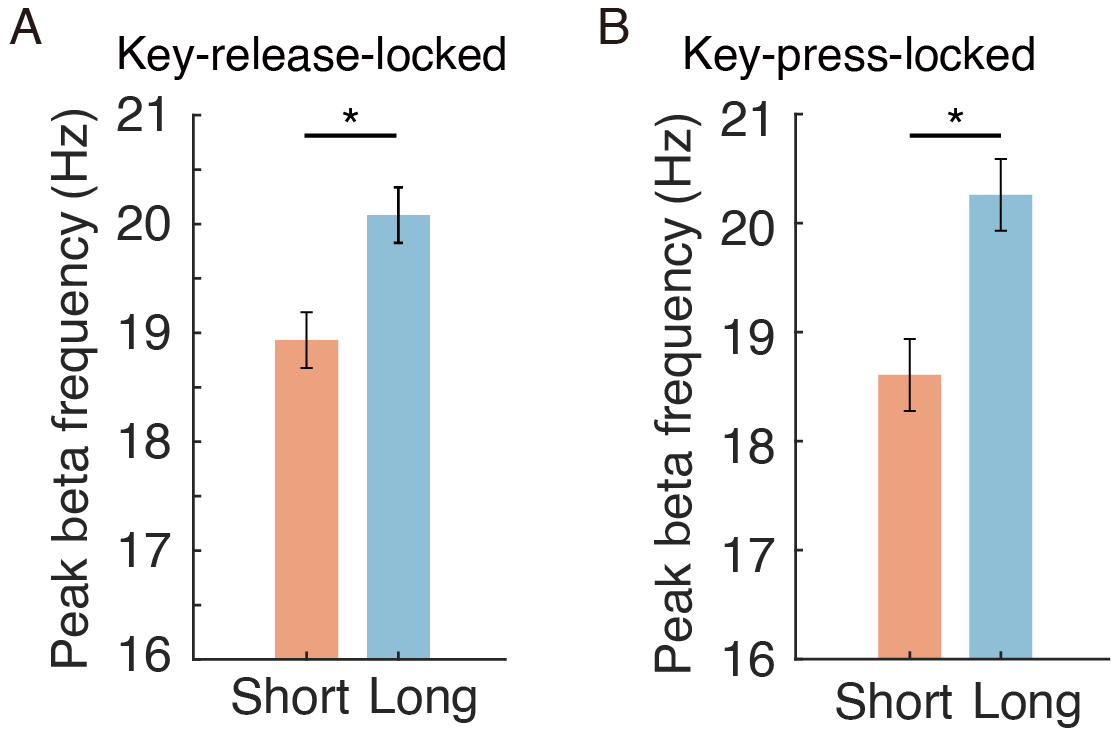

**Figure S3. Relationship between peak beta frequency and self-timed duration. A**, Peak beta frequency (PBF), averaged across all electrodes and participants, for the long- and short-duration conditions in the key-release-locked analysis, computed over the pre-release period (-2 to 0 s relative to key release). **B**, Same as (A), but for the keypress-locked analysis, with PBF computed within the -0.75 to 0.75 s window around keypress onset. Error bars indicate ±1 within-subject SEM. Asterisks denote significant differences between conditions (* *p* < 0.05).

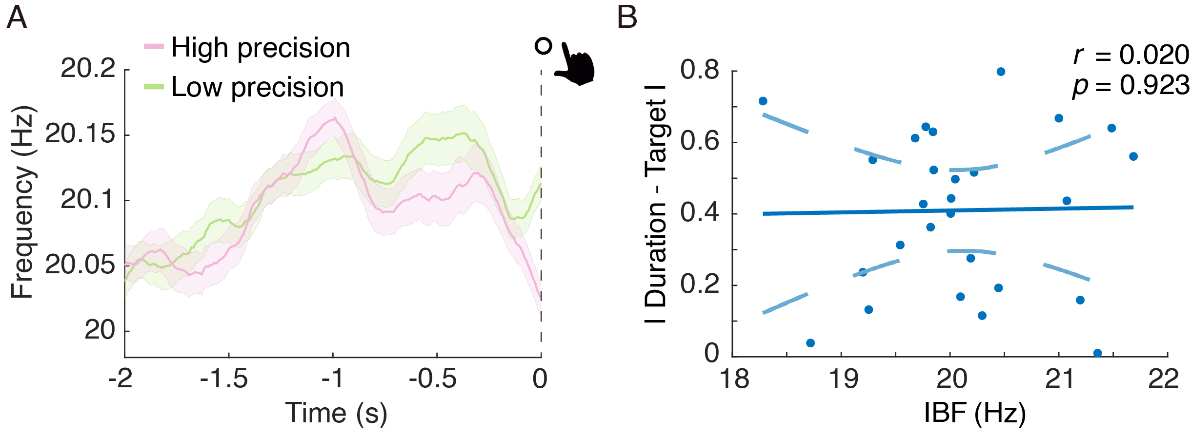

**Figure S4. Relationship between instantaneous beta frequency and absolute deviation from the target duration. A**, Instantaneous beta frequency for high-precision and low-precision trials in the key-release-locked analysis. Precision was defined based on the absolute deviation of self-timed duration from the target duration (1.5 s): trials in the lower tertile (smaller absolute error) were classified as high-precision trials, whereas trials in the upper tertile (larger absolute error) were classified as low-precision trials. No significant differences were observed between conditions (*cluster-based correction*, all *p*s > 0.05). Shaded regions denote ±1 within-subject SEM. **B**, Across-participant relationship between mean instantaneous beta frequency (IBF), averaged across all electrodes, and the absolute deviation from the target duration (1.5 s). No significant correlation was observed (*r* = 0.020, *p* = 0.923).

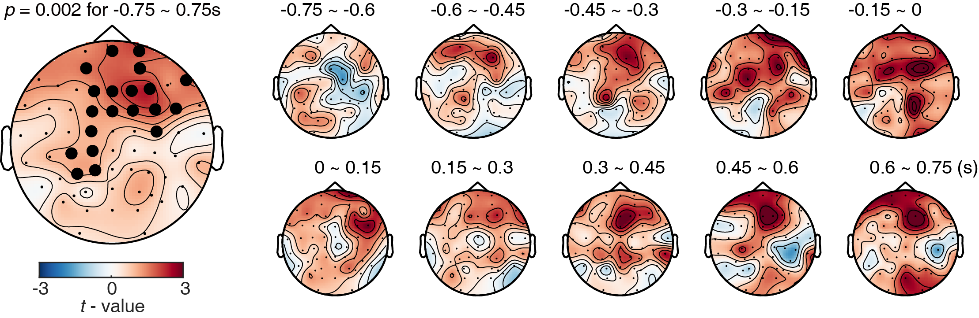

**Figure S5. Spatial characterization of beta-frequency modulation in key-press-locked analysis.** Left: scalp topography of beta-frequency modulation associated with self-timed duration. Beta-frequency values were averaged for each electrode within the keypress-centered window (-0.75 to 0.75 s relative to keypress onset) for the long- and short-duration conditions. Electrodes showing significant differences (long > short) are indicated by bold circular symbols (*cluster-based correction*, *p* = 0.002). Right: dynamic scalp maps showing beta-frequency differences (long-short) across consecutive 0.15-s intervals within the keypress-centered period, illustrating how the modulation evolved over time.

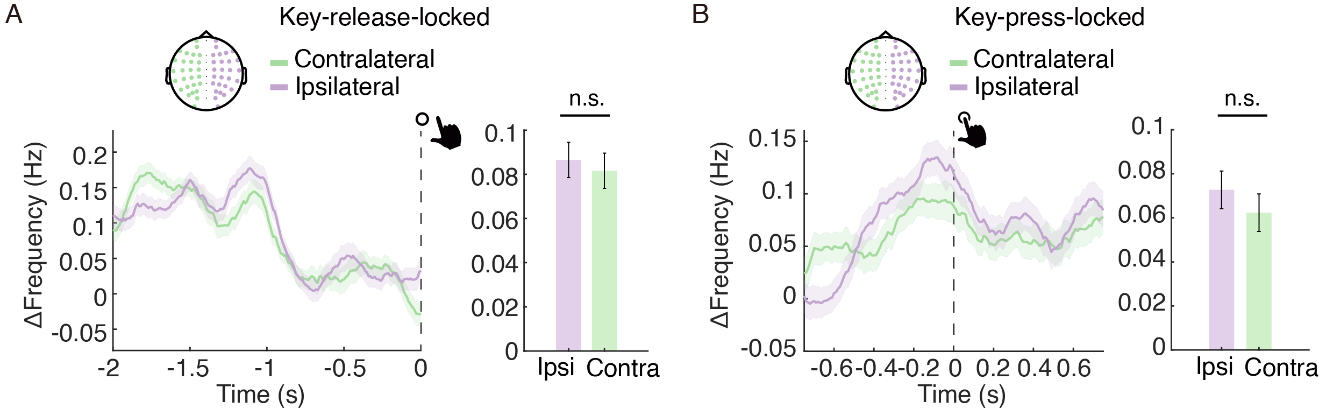

**Figure S6. Comparison of beta-frequency modulation between contralateral and ipsilateral electrode sites relative to the hand used. A,** The left panel shows the time course of beta-frequency modulation across the pre-release period (-2 to 0 s relative to key release); no significant ipsilateral-contralateral differences were observed (*cluster-based correction,* all *p*s > 0.1). The right panel shows the mean beta-frequency modulation averaged over this period for the two conditions; the ipsilateral and contralateral averages did not differ significantly (*t_26_*= 0.305, *p* = 0.763). **B,** Same as (A), but for the keypress-locked analysis. The left panel shows the time course across the keypress-centered window (-0.75 to 0.75 s relative to keypress onset); again, no significant ipsilateral-contralateral differences were detected (*cluster-based correction,* all *p*s > 0.1). The right panel shows the corresponding mean beta-frequency modulation for the two conditions, which also did not differ significantly (*t_26_*= 0.610, *p* = 0.547).

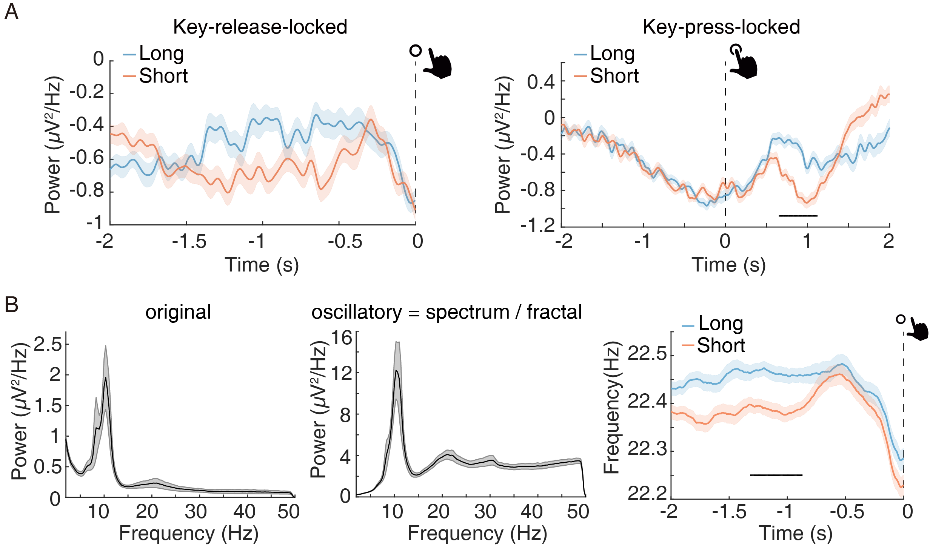

**Figure S7. Control analyses excluding power and spectral contributions to beta-frequency modulation. A**, Beta power for the long- and short-duration conditions averaged over all channels in the key-release-locked analysis (Left) and the key-press-locked analysis (right). **B,** The original amplitude spectrum (left) and the oscillatory component of the spectrum (middle) obtained by separating the aperiodic component. Instantaneous beta frequency (right) for the long- and short-duration conditions averaged over all channels in the key-release-locked analysis after removing 1/f component. Significant time points are indicated by the horizontal black bar (*cluster-based correction*, *p* < 0.05). Shaded regions denote ±1 within-subjects SEM.
